# Characterizing Oligodendrocyte-Lineage Cells and Myelination in the Basolateral Amygdala: Insights from a Novel Methodology in Postmortem Human Brain

**DOI:** 10.1101/2025.01.09.631560

**Authors:** Kelly Perlman, Emanuel Curto, Sarah Barnett-Burns, Maria-Antonietta Davoli, Volodymyr Yerko, Reem Merza, Andreas I Papadakis, Alan Spatz, Christopher R Pryce, Gustavo Turecki, Naguib Mechawar

## Abstract

The basolateral amygdala (BLA) plays a key role in the pathophysiology of depressive disorders and trauma, yet oligodendrocyte-lineage cells and myelin in this brain region remain understudied in humans. This may be due, at least in part, to the lack of a cost-effective, antibody-based method to isolate oligodendrocytes (OL) and OL precursor cells (OPC) from postmortem brain tissue that is compatible with molecular biology applications. This study aimed to 1) create and validate a method for isolating OPC and OL nuclei from frozen postmortem grey matter; 2) compare OPC and OL gene expression in the BLA between individuals with depression who died by suicide (with or without a history of childhood abuse) and matched controls; and 3) provide histological characterizations of OPC, OL, and myelin in the BLA. Frozen left-hemisphere BLA samples were obtained from brain donors with well-characterized phenotypic information. Immunolabeled nuclei were sorted into OPC (SOX10+/CRYAB-) and OL (SOX10+/CRYAB+) populations, and RNA was measured using a custom Nanostring codeset. Fluorescence in situ hybridization was used to determine OPC (PDGFRα+) and OL (MYRF+) densities, and immunofluorescence was used to label axons (NF-H) and myelin (MBP) for myelin area fraction. The method successfully isolated OPC and OL nuclei with correct transcriptomic profiles. In the OL fraction, MOBP was found to be significantly decreased in depressed individuals with a history of child abuse compared to controls, but no other genes showed significant group differences in either fraction. However, significant age-related patterns were observed in both fractions. Furthermore, no significant differences in cell densities or myelin coverage were observed across groups. Lastly, a strong significant correlation between OL density and myelin area fraction was identified. This study provides a novel sorting method and a comprehensive characterization of OL-lineage gene expression, cell densities, and myelin in the human BLA.

## 1. Introduction

Oligodendrocytes (OLs) extend processes that ensheathe axons in the central nervous system, forming myelin, which provides electrochemical insulation that increases the conduction velocity of action potentials (Waxman, 1980). Myelination is an activity-dependent phenomenon, as observed following environmental changes such as social isolation or with the acquisition of new skills (Williamson and Lyons, 2018). Changes in myelination that alter conduction velocity and timing of information transmission are a form of life-long plasticity (Fields, 2015).

The importance of gray matter OL-lineage cells and myelin in brain health and disease is becoming increasingly evident (Knowles et al., 2022; Spieth and Simons, 2024; Timmler and Simons, 2019; Werkman et al., 2021). Although the majority of studies in the field have been conducted in rodents, recent evidence has highlighted major differences between mouse and human OL gene expression and myelin composition (Gargareta et al., 2022). Such observations further highlight the need for more studies conducted in humans. For example, there is currently a paucity of region-specific characterization of OL-lineage cells and myelination in the postmortem human brain with respect to gene expression profiles and histological properties, among other metrics. Single nucleus RNA sequencing (snRNAseq) technologies are pushing this field forward, characterizing different subpopulations of OLs and oligodendrocyte precursor cells (OPC) (Jäkel et al., 2019; Maitra et al., 2023; Van Bruggen et al., 2017). However, this technology can still be prohibitively expensive and therefore out of reach to a significant part of the research community. It would be beneficial to have a protocol to isolate OPC and OL nuclei from frozen archived postmortem human brain tissue that would be reliable, cost-effective, and compatible with targeted downstream experiments to investigate the healthy and diseased brain.

An interesting body of literature is emerging which highlights depression-associated dysregulation in OL/myelin in humans as well as related animal models (Sharma et al., 2023; Zhou et al., 2021). In depression, region-specific compromised myelin integrity, morphological and density differences in OL-lineage cells, as well as changes in OPC and OL gene expression, have been reported (Aston et al., 2005; Boda, 2021; Nagy et al., 2020; Sibille et al., 2009; Zhou et al., 2021). Individuals with a history of childhood abuse (CA) who suffer from depression have a more severe course of illness and have higher rates of treatment resistance, possibly constituting a unique neurobiological subtype (Teicher and Samson, 2013). Our group has previously identified CA-specific changes in gene expression and methylation converging on OL/myelin pathways, as well as thinner myelin sheath in small caliber axons in the anterior cingulate cortex (Lutz et al., 2017). Other limbic regions, such as the amygdala, are also of obvious interest but have yet to be explored in this regard.

The amygdala is a mid-temporal lobe structure central to the generation of emotional responses, and learning and consolidation of emotionally-valanced memories (Janak and Tye, 2015). Though the precise neuroanatomical groupings of its subdivisions are still debated, the amygdala contains three principal groups: the basolateral (BLA), the centromedial, and the cortical-like amygdalar groups (Yang and Wang, 2017). The amygdala is a key hub in major circuits that control affective behavior and mood, and it displays aberrant activity in various mental health conditions, including in depressed patients and in individuals with a history of CA (Gee et al., 2013; Ho et al., 2022; Teicher et al., 2016; Zhu et al., 2019). As such, it is crucial to have a baseline characterization of human OL-lineage and myelin properties in the human amygdala to understand how it might be altered in stress-related psychiatric disorders. Understanding how OL and myelin level differences contribute to altered activity in the amygdala is necessary for a comprehensive characterisation of a depression or CA neurophenotype.

A recent region-specific study of chronic social stress, a common mouse model of depressive-like behavior, revealed downregulation of numerous myelin genes (e.g., Mog, Opalin, Mobp, Mag, Mbp, Plp1, Cnp) in the BLA of adult male mice via RNA sequencing (Cathomas et al., 2019). In the current study, we aimed to observe whether this striking pattern in mice would be mirrored in the BLA of humans with depression. Specifically, we wanted to understand whether a downregulation of myelin-constituent genes in the BLA is part of the underlying neurobiology of depression, or whether it reflects a more general stress-specific adaptation, as has been observed in other brain regions in individuals with a history of CA (Teicher et al., 2016). This study was thus conceptualized with three complementary aims: First, to create and validate a novel sorting method to isolate frozen grey matter nuclei into OPC and OL populations. Second, to apply this new method to characterize the OPC- and OL-specific gene expression profiles of individuals with depression who died by suicide with or without a history of severe CA. Finally, we aimed to determine OPC and OL cell densities, as well as the fraction of axon area covered by myelin, and to compare these metrics between groups. By combining these complementary approaches, our overall aim was to achieve a comprehensive understanding of baseline profiles of OL-lineage cells and myelin in the human BLA and how they might be affected in depression and/or by a history of CA.

## 2. Methods

### 2.1 Post-Mortem Human Brain Samples

The Douglas Hospital Research Ethics Board approved this study. Informed consent was received from the next of kin of each brain donor. Fresh-frozen BLA samples were provided by the Douglas-Brain Canada Brain Bank (www.douglasbrainbank.ca). Relevant donor information was obtained through the standardized psychological autopsy way of proxy-based interviews with next of kin (adapted from SCID-I and SCID-II), as described previously (Dumais et al., 2005), combined with information provided by the Quebec Coroner’s office, as well as medical and social services records, when available. Childhood maltreatment was assessed by way of an adapted Child Experiences of Care and Abuse Interview (Bifulco et al., 1994). Toxicology was performed to assess the presence of substances in the blood at time of death.

Tissue samples were matched between three groups: 1) control (CTRL) individuals having died suddenly (i.e., from accidents or physical illness) in absence of psychiatric illness and with no history of CA; 2) patients with depression who died by suicide and had no history of CA (DS); 3) patients with depression who died by suicide and who had a history of severe CA (DS-CA). Neurodegenerative disorders were exclusion criteria for all groups.

Dissections were performed by expert brain bank staff with the guidance of a human brain atlas (Mai et al., 2015). Whole brain samples were initially cut into serial 1 cm-thick slabs. The ventromedial third of the left amygdala was isolated to extract the BLA. Samples were stored unfixed at -80 °C until further processing.

### 2.2 Nuclei preparation and sorting

Frozen BLA (less than 100 mg of tissue) was homogenized in a glass douncer in 3 ml of 0.1% tween lysis buffer with RNAseInhibitor (Sigma, cat# 3335402001, 1:1000). After 4 min, 10 ml of 5% bovine serum albumin (BSA) wash buffer was added, the suspension was filtered through a 30 µm MACS filter, and then was spun at 3500 g for 6 min at 4°C. The pellet was then resuspended in 5 mL of 2% BSA wash buffer, filtered through a 30 µm MACS filter, and spun again at 3500 g for 6 minutes at 4°C. After resuspension in 400 µl of PBS, the nuclei suspension was incubated for 1 h in 100 µl of 5% BSA blocking buffer (with normal donkey serum - NDS), Goat anti-Sox10 (R&D Systems, cat# AF2864, 1:100), Mouse anti-CRYAB (Origene, cat# TA500583, 1:100), and RNAseInhibitor (1:500) at 4°C on a rotor. The nuclei were then spun at 500g for 3 min at 4°C. After resuspension in 400 µL 1x PBS, the nuclei suspension was incubated for 1 h in 100 µl of 5% BSA blocking buffer (with NDS), Donkey anti-Goat Alexa 488 (1:500), Donkey Anti-Mouse Alexa 647 (1:250), and RNaseinhibitor (1:500). Finally, the nuclei were spun at 500 g for 3 min at 4°C, resuspended in 500 µl of PBS with Hoescht dye (1:500) and RNaseInhibitor (1:500), and kept on ice until sorting. Nuclei were sorted at 4°C on FACSAria Fusion (BD Biosciences). The populations were gated by first isolating singlets using the particle size (forward scatter) and Hoescht signal (375-nm laser, 450/50 bandpass filter, 410 longpass filter), then by isolating the SOX10+ population using SOX10/Alexa488 immunofluorescence (488-nm laser, 530/30 bandpass filter, 502 longpass filter), and finally by splitting into OPC- and OL-enriched fractions by CRYAB/Alexa647 immunofluorescence (640-nm laser, 670/30 filter). We deemed the SOX10+/CRYAB-negative population as OPC-enriched (termed OPCs) and the SOX10+/CRYAB+ fraction as OL-enriched (termed OLs). See Figure 2a for a gating example.

**Figure 1.**
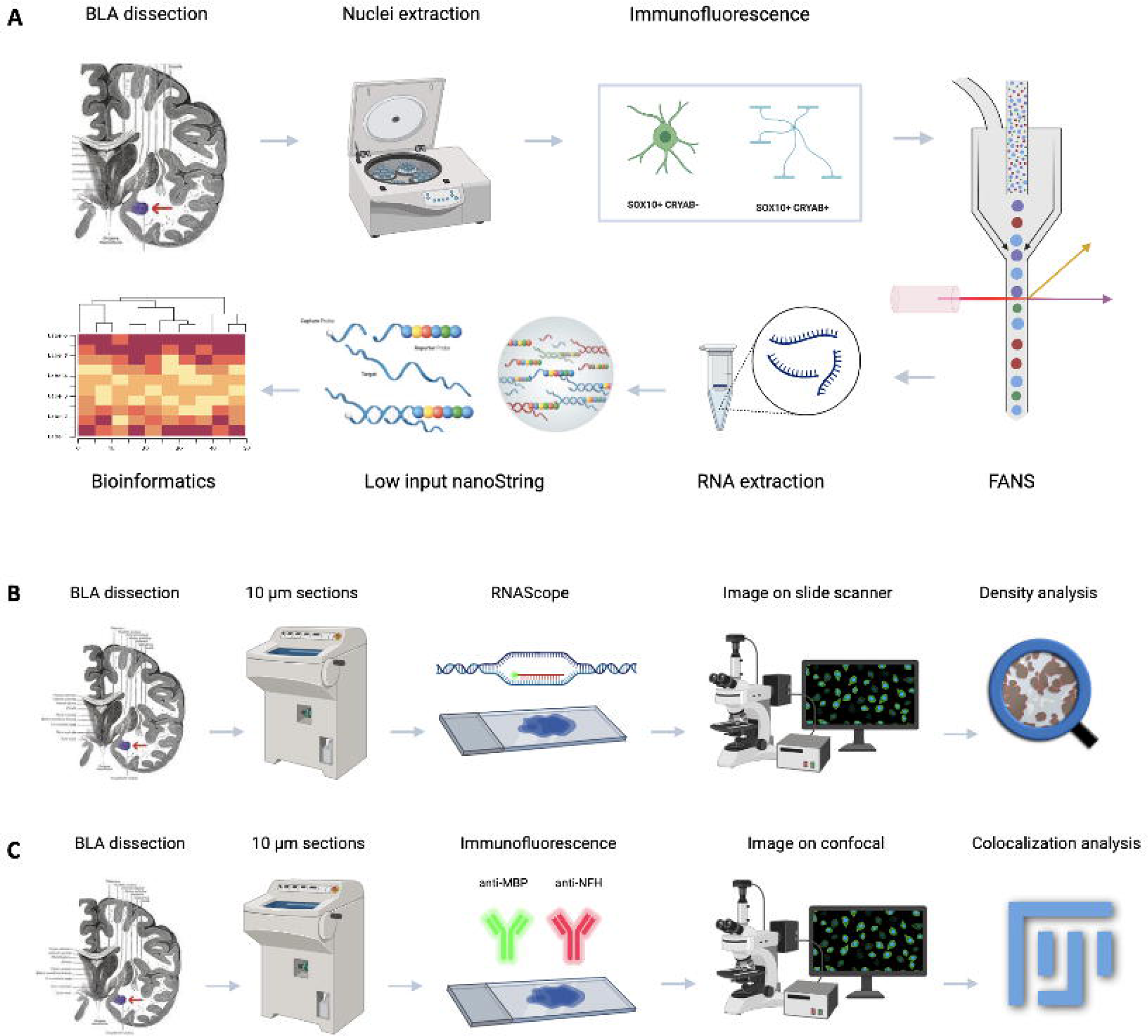
Graphical overview of methods. A) Nuclei preparation and FANS sorting method to generate OPC and OL populations, which are then used as input to a targeted Nanostring panel. B) RNAscope-based density measurements for OPC and OL. C) IF-based myelin area fraction coverage of axons. FANS: fluorescence-assisted nuclear sorting, IF: immunofluorescence, OPC: oligodendrocyte precursor cells. OL: oligodendrocytes. Figure made with biorender.com

**Figure 2.**
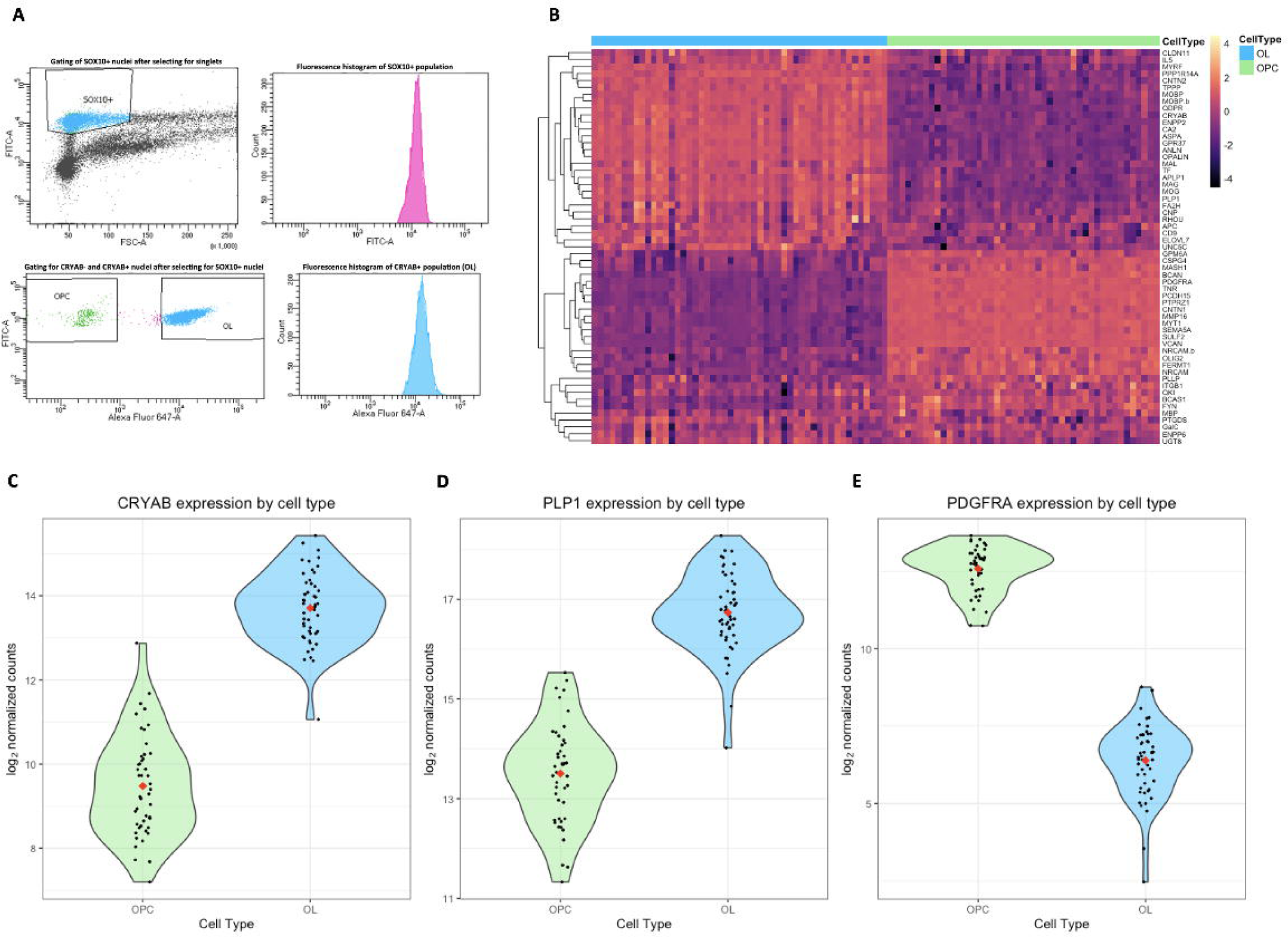
Validation of novel OL-lineage nuclei sorting method. A) Representative FANS plot demonstrating population gating by SOX10 and CRYAB antibodies. Note that the “intermediate” cells between the OPC and OL gating on the SOX10 vs CRYAB plot are not collected, in order to avoid possible contamination, given the identity of these cells on the OL-lineage is unclear. B) Heatmap demonstrating the gene expression for all subjects, where lighter colors represent higher z-scored expression and darker colors represent lower z-scored expression. The annotation bars for each column on the heatmap cell type information (OPC in green, OL in blue). Violin plots showing gene expression between OPC and OL cell types of C) CRYAB, D) PLP1, and E) PDGFRα. The red diamond represents the mean expression per cell type population. FANS: fluorescence-assisted nuclear sorting, CTRL: control, DS: depressed suicides, DS-CA: depressed suicides with a history of child abuse. OPC: oligodendrocyte precursor cells. OL: oligodendrocytes.

Note that on the forward scatter vs. SOX10+ fluorescence plot, the larger particles (generally above 150 x 10^3^ on the FSC axis) were excluded. OL-lineage nuclei are typically quite small (Wolswijk, 2000), and these larger particles are likely nuclei with debris attached to them, which presumably contains ambient RNA, thus confounding the signal. SOX10 is a well-known transcription factor that is highly specific for OL-lineage (Kuhlbrodt et al., 1998). CRYAB was selected because it is expressed in several cellular compartments, including the nucleus (Van Rijk et al., 2003) and it is highly expressed in OL but not OPC (Perlman et al., 2020). The combination of these two markers allowed us to gate the flow cytometry properly to specifically isolate the nuclei of OPC and OL. We sorted on “purity” mode at speed 1.0 on an 85 µm nozzle on the BD FACSAria.

### 2.3 RNA extraction and nanoString panel

OPC and OL nuclei were each sorted directly into Buffer RL from the Single Cell RNA Purification Kit (Norgen) and total RNA was extracted with this same kit. Concentrations were quantified with High Sensitivity RNA TapeStation (Agilent). The custom Nanostring codeset was designed to work with nCounter® Low RNA Input Kit.

The codeset was created to be exon-spanning, in order avoid having to deplete for genomic DNA, as in our experimental experience, the depletion steps tend to be too harsh on small, sorted populations (e.g., OPC) rendering them unusable for downstream applications. Even though more than half of the mRNA in the nucleus is unspliced, it still contains a substantial amount of spliced mRNA (Bakken et al., 2018). Therefore, all probes were created to be exon spanning, except for CLDN11, which was not computationally possible according to Nanostring bioinformaticians.

This codeset was designed for maximum isoform coverage, so for 2 genes (MOBP and NRCAM) it was necessary to make 2 probes to cover all key isoforms. The codeset also contains 6 housekeeping genes for normalization (AARS, ABCF1, MTO1, POLR1B, SDHA, TBP).

Nanostring results have been previously found to be almost perfectly correlated with RNA sequencing results (Lutz et al., 2017, supplemental material). The genes in the codeset were selected via literature review (Cahoy et al., 2008; Lutz et al., 2017) and the full codeset design, including all the genes and captured isoforms, can be found in Supplementary File 1. The aim was to focus on OL-lineae and myelin specific targets, to be able to compare with existing literature. The nCounter Low RNA Input workflow was performed at the Nanostring Gene Expression Profiling Core Facility of the Lady Davis Institute. Figure 1a shows a graphical summary of this methodology.

The Nanostring raw data (.RCC files) was processed with nSolver Analysis Software version 4.0. Genes were removed from the analysis if they had a value for “percent samples above threshold” under 60%, which eliminated NEU4, GSX2, TMEM88b. However, most of the genes were above the detection threshold in over 90% of the samples. Samples were removed from the analysis if they had a value for “percent probes above threshold” under 75%.

Background thresholding was performed by taking the mean plus 2 standard deviations of the counts for the negative control probes, as recommended by Nanostring (https://nanostring.com/wp-content/uploads/Gene_Expression_Data_Analysis_Guidelines.pdf). This takes any gene counts lower than the designated threshold and floors them to that value, preventing inflated fold changes for low expressing genes. Positive control normalization was performed to correct for platform-specific variability, and housekeeping gene normalization was performed to correct for sample input variability. Normalization factors for each lane were calculated in nSolver (with geometric means selected for both the positive controls and the housekeeping genes). The gene counts are shown as log_2_ normalized gene expression.

### 2.4 RNAScope fluorescence in situ hybridization

Frozen-unfixed BLA blocks stored at -80°C were cut using a Leica CM1950 cryostat. The tissue was sectioned from the rostral side at 10 µm and collected onto Superfrost-charged slides. Slides were stored at -80°C until further processing. RNAScope® probes and reagents were acquired from Advanced Cell Diagnostic; the manufacturer’s protocol was followed throughout the *in situ* hybridization. First, slides were placed in 4°C, 10% neutral buffered formalin for 15 min to fix the tissue. Five successive dehydrations (at 50%, 70%, 95%, 100%, 100% ethanol concentrations) occurred, each lasting 2 minutes. Slides were air dried for 5 min. To quench endogenous peroxidase activity, 3% hydrogen peroxide reagent was applied to the samples for 10 min at room temperature. Protease digestion was allowed to occur for 30 min at room temperature, to permeabilize cells. In a 40°C humidity-controlled oven (HybEZ II, ACDbio), the following probes were hybridised for 2 hours: Hs-PDGFRα and Hs-MYRF. The addition of amplifiers was achieved by proprietary AMP reagents; signal visualisation took place by probe-specific horseradish peroxidase-based detection and tyramide signal amplification using Opal dyes (Opal 520 – 1:700, Opal 570 – 1:500). True Black was used to reduce background and autofluorescence during imaging: True Black (1:20) in 70% ethanol was applied to tissue for 45 sec; subsequently, the tissue was washed thoroughly in dH_2_O. Mounting medium 4′,6-diami-dino-2-phenylindole (DAPI) for nuclear staining (Vector Laboratories) was used to coverslip the samples, which were subsequently stored at 4°C until imaging. Figure 1b shows a graphical summary of this methodology.

### 2.5 Immunofluorescence

Slides with frozen-unfixed BLA samples, sectioned and stored as described above, were fixed in 10% neutral buffered formalin for 5 min at room temperature and then rinsed in PBS. Samples were blocked in 5% NDS diluted in 0.2% Triton X-100 in PBS for 1 h. Samples were then incubated overnight with primary antibodies chicken anti-NF-H (abcam, cat# ab72996, 1:1000) and mouse anti-MBP (BioLegend, cat# 80840, 1:3500) diluted in PBS-Tx-100 with 5% NDS. They were then rinsed in PBS and incubated for 1 h with HRP conjugated secondary antibodies: donkey anti-chicken (1:500) for NF-H and donkey anti-mouse (1:500) for MBP, both diluted in PBS-Tx-100 with 5% NDS. To reduce background and autofluorescence during imaging, samples were treated with True Black (1:20) for 20 sec and washed thoroughly with dH2O. Samples were coverslipped with DAPI containing mounting medium (Vector Laboratories). Figure 1c provides an illustrated summary of this methodology.

### 2.6 OPC and OL density measurements

Slides were imaged using an Olympus VS120 Slide Scanner at 20x magnification. Images were processed by a blinded researcher on QuPath (v 0.5.1). Cells with three or more puncta of PDGFRα co-localised with a DAPI nucleus, without visible MYRF puncta, were counted as OPCs. Alternatively, three or more MYRF puncta co-localised with DAPI signal were counted as OLs. Six regions of interest (ROI) of equal size were chosen per sample; ROIs were chosen at spatially distinct, non-overlapping parts of the image based on the absence of tissue folds, holes, dust, or other artefacts. Average density was determined for each subject as total cell counts divided by the total area of the six ROIs.

### 2.7 Myelin Coverage and Area Fraction measurements

Images were taken on a FV1200 Olympus confocal microscope at 40x magnification with z-stack steps of 0.5 µm. At least 3 images from each given section were taken at spatially distinct locations. Images were processed using Fiji version 2.0.0. The maximum intensity projection was applied to each composite z-stack image. Each image was split into its respective channel and the MBP and NF-H channels were thresholded individually, using the Otsu automatic segmentation method (Otsu, 1979). Subsequently, a binary mask was created for each channel.

The binary masks were input to JACoP: Just Another Colocalization Plugin (Bolte and Cordelières, 2006), and the Mander’s M2 coefficient was extracted for each image (Costes et al., 2004; Manders et al., 1993). The M2 coefficient represents the ratio of the MBP signal 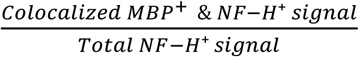. The multiple images taken per subject were averaged to provide a single M2 value for each subject. Frames were removed if M2 signal > 0.5, as this would not be realistic for gray matter, and/or if MBP signal appeared non-specific. For most of the M2 signals > 0.5, MBP signal appeared non-specific.

### 2.8 Statistics

All statistics were conducted with R version 4.4.3. To determine whether age, PMI, and pH covariates were statistically different between groups, one-way anovas were used. For all group comparisons, an ANCOVA was run for each metric (OPC density, OL density, ratio of OL/OPC), with age, sex, and pH as covariates, such that the effect of group could be isolated from potential confounders. PMI was excluded as a covariate after it was found that removing it consistently improved model fit as determined by Bayesian Information Criterion (BIC). To determine the best model to fit the relationship between age and gene expression, we made linear, quadratic, cubic, and logarithmic models for each gene, and then obtained the BIC) as a measure of model fit. Generally, the lowest BIC value among all the models tested indicates the best fitting model. However, an absolute BIC value difference of less than 2 does not provide sufficient evidence for one model over another (Berchtold, 2010). Therefore, we considered the linear model as the “default” due to ease of interpretability, unless the BIC for the quadratic, cubic, or logarithmic models was less than the linear model BIC by at least 2. In other words, if BIC_linear_ – BIC_other_ ≥ 2, the non-linear model was chosen, as it indicates that the non-linear is an improvement on model fit. Pearson’s correlations were used to measure correlations between M2 coefficient and cell densities. P < 0.05 was used as the significance threshold for all statistical analyses. Data are represented as mean ± standard error of the mean.

## 3. Results

### 3.1 OPC- and OL-specific expression of target genes

Figure 2a illustrates an example FANS plot with gating of OPC and OL populations based on fluorescence associated with SOX10 and CRYAB antibodies. The OPC and OL fractions for each subject were each represented in the Nanostring panel, except for a few subjects that were excluded from each fraction due to quality control concerns (see Methods) or subjects that had insufficient OPCs to generate reliable results. As such, Supplementary Tables 1 and 2 show the subject information separately for all subjects that had usable data for OPC fraction and OL fraction, respectively.

To ensure that white matter contamination or differences in SOX10+ gating were not driving any group differences, we compared the average percentage of SOX10+ singlets represented in the FANS plots across groups (Supplementary Figure 1). The average SOX10+ percentage across all sorts was 31.18%, and we observed no statistically significant differences between groups, indicating that it would be unlikely for any group result to be driven by contamination or gating inconsistencies.

Principal component analysis (PCA) revealed that the OPC and OL fractions were clearly separated across principal component 1, which explained 75.9% of the variance in the data (Supplementary Figure 3). Gene expression of OPCs and OLs matched the expected patterns based on the literature, with the OPC fraction enriched for PDGFR𝛼, CSPG4 and PTPRZ1, while the OL fraction was enriched for MOG, MAG, and PLP1 (Figure 2b). To validate the gating, we saw that, as expected, OPCs showed low levels of CRYAB expression while OLs showed far higher expression (Figure 2c). We observed a similar pattern for PLP1, a canonical mature OL marker that codes for the most abundant protein in the myelin sheath. It is worth noting that while PLP1 expression is far lower in OPCs compared to OLs, it is still expressed consistently in the former (Figure 2d). Together, this suggests that the OPCs captured here are OPCs that are “later” along the continuum of OPC maturity, consistent with what has been observed in adults (Perlman et al., 2020). However, they can nonetheless be designated as progenitors through their expression of canonical markers like PDGFRα (Figure 2e). Notably, the violin plots in this figure also show high variability in gene expression, consistent with what is observed typically in postmortem human brain samples. Altogether, these results validate our novel nuclei sorting method, indicating that the correct cell types were isolated, with RNA that was usable for quantitative measurements.

In the OPC fraction, we did not observe any genes that showed statistically significant differences in gene expression between groups. In the OL fraction, we found that MOBP was significantly different between groups (p = 0.033), specifically lower in the DS-CA group compared to CTRL. This group difference remained significant after correction for multiple comparisons using the Benjamini-Hochberg method within each model (p = 0.044) (Supplementary Figure 4A). Two MOBP probes (MOBP and MOBP.b) had to be produced to cover all key isoforms, and while probes both showed the same pattern across groups, MOBP.b was not statistically significant (p = 0.20) (Supplementary Figure 4B). No other genes were found to be significantly differently expressed across groups in the OL fraction. Taken together, these findings show that in the OPC and OL fractions, there is no robust effect of depression or CA in the left BLA with respect to expression of these specific target genes.

Given that the subjects in this study had an age range between 19 and 73 years, we plotted gene expression patterns across a wide age span and modelled gene expression as a function of age. We elected to remove the group term from the model as the average BIC across models was lower when removing this term, indicating better model fit (mean BIC of OPC and OL models decreased by 6.18 and 6.53, respectively when removing the group term). In both the OPC and OL fractions, all significant models displayed linear relationships. In the OPCs, BCAS1, CNTN1, ENPP6, FYN, MOG, TF, and UGT8 showed decreases across the age span, while IL5, ITGB1, MASH1, PCDH15, PPP1R14A, and TPPP showed increases across the age span (Figure 3a). In the OL fraction, the expression of most genes was decreased with age (ENPP6, GalC, TF, UGT8), though BCAN was increased with age (Figure 3b). The expression of ENPP6, TF, and UGT8 were decreased with age in both fractions. In summary, the age relationships between genes in each fraction were distinct, but with some overlap.

**Figure 3.**
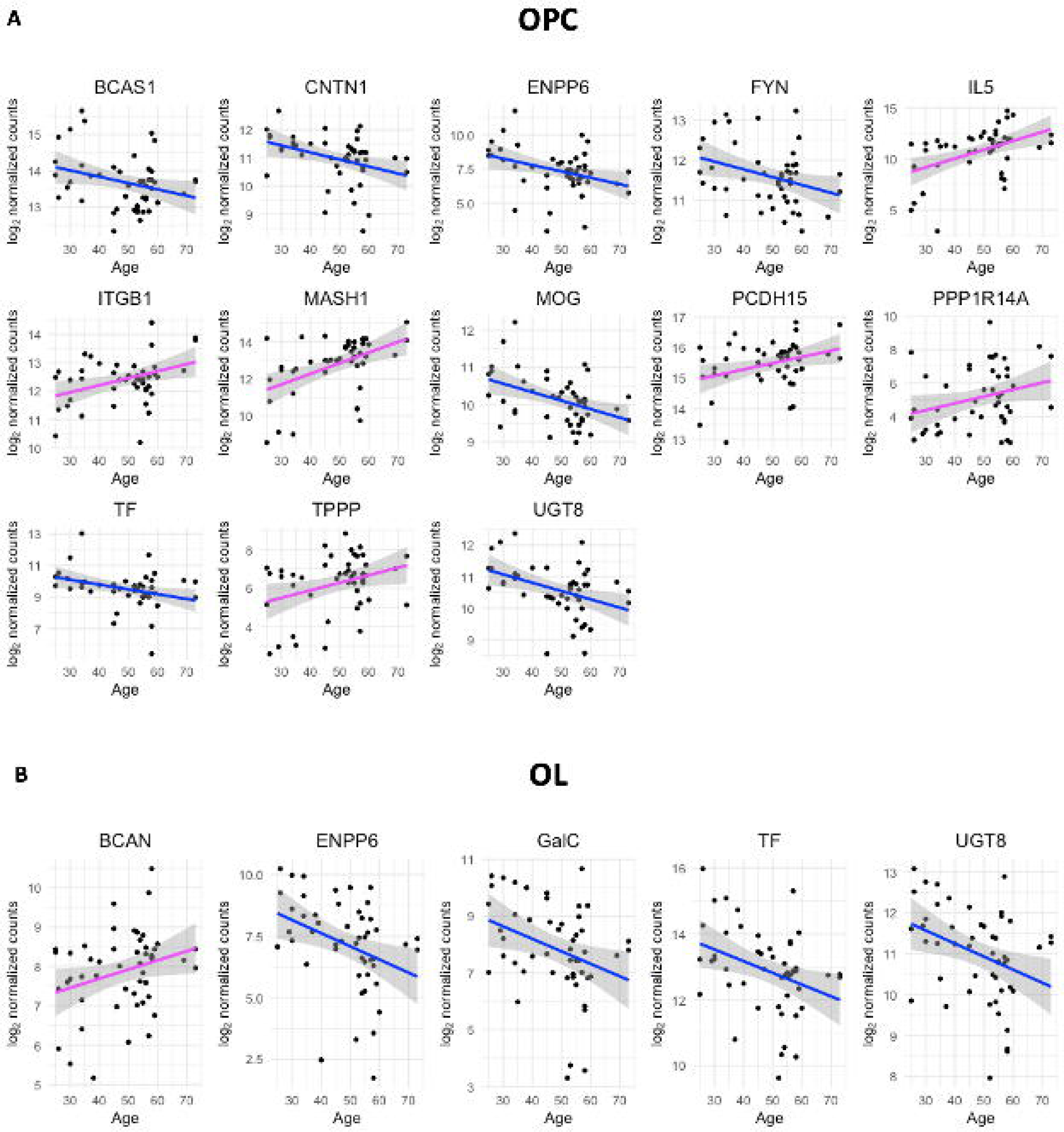
OPC- and OL-specific gene expression as a function of age. Scatter plots with smoothed regression lines for each gene that shows a statistically significant relationship with age in both A) OPC and B) OLs. If the model has a negative coefficient for the age term, the regression line is blue and if the model has a positive coefficient for the age term, the regression line is magenta. All significant OPC and OL gene models show linear relationships. OPC: oligodendrocyte precursor cells. OL: oligodendrocytes.

### 3.2 OPC and OL Density

OPCs were visualised by PDGFRα and DAPI colocalization, and OLs were visualised by MYRF and DAPI colocalization (Fig. 4a). Subject information for the density analyses can be found in Supplementary Table 3. The overall average OPC density was 31.61 ± 1.40 cells/mm^2^, the average overall OL density was 111.31 ± 13.77 cells/mm^2^, and the average OL/OPC density ratio was 3.78 ± 0.511. The effect of group did not reach statistical significance in any model (OPC density: p = 0.12, OL density: p = 0.33, OL/OPC ratio: 0.50). None of the covariates in any of the models reached statistical significance except for sex (p = 0.0058), in which mean OPC density was higher in males (33.8 ± 0.94 cells/mm^2^) compared to females (25.7 ± 3.86 cells/mm^2^) (Supplementary Figure 5). However, when observing the plots descriptively, OPC density was slightly higher in DS-CA compared to DS and CTRL groups, but more strikingly OL density appeared to be lower in DS-CA and DS groups compared to controls, and correspondingly OL/OPC ratio followed the same pattern (Figure 4b). There was no statistically significant relationship between age and OPC density (Supplementary Figure 6a, p = 0.49), age and OL density (Supplementary Fig 6b, p=0.38), or age and OL/OPC ratio (Supplementary Figure 6c, p = 0.67).

**Figure 4.**
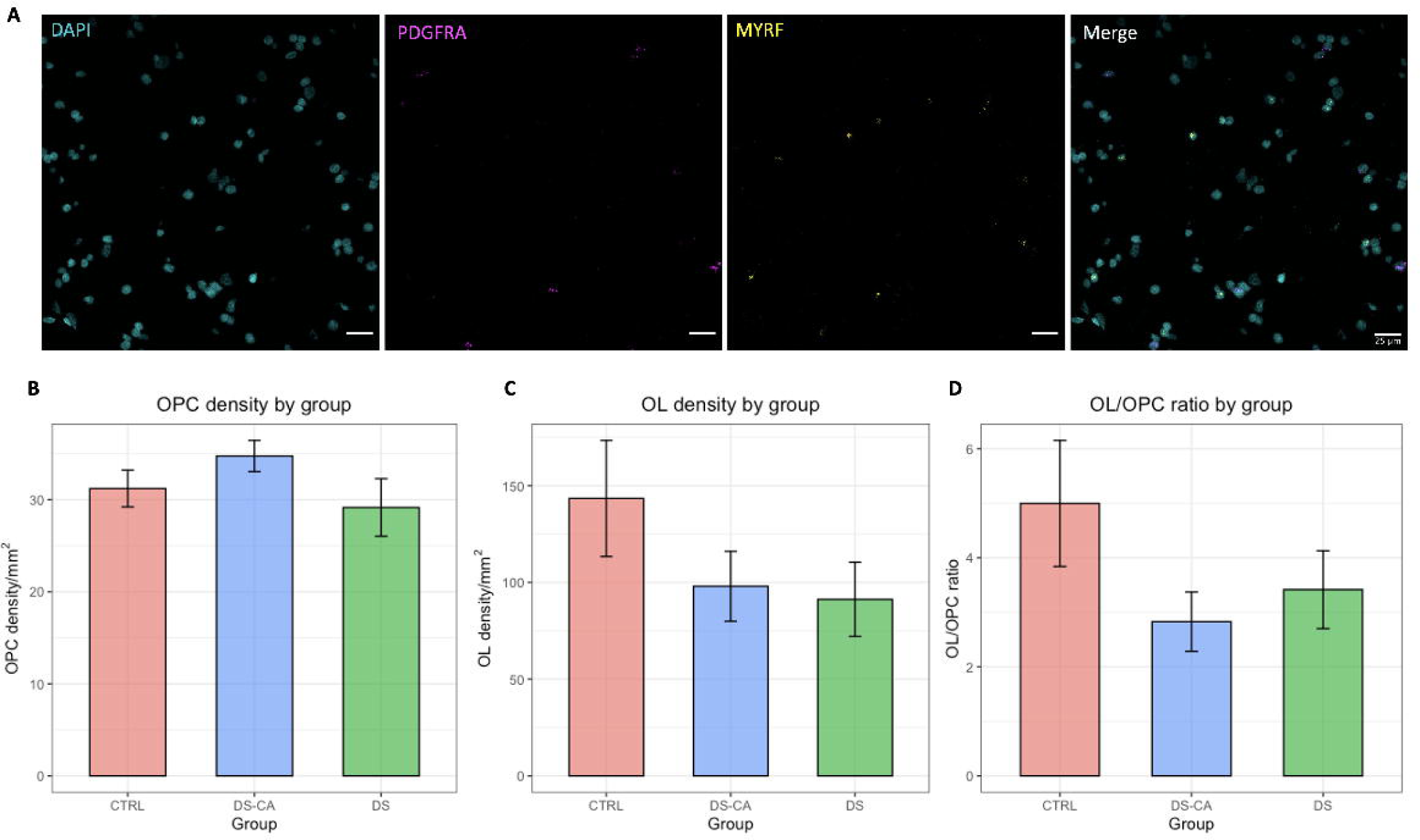
OPC and OL density analysis. A) Representative Images of RNA in situ hybridization. Left: DAPI staining of the nucleus (cyan); PDGFRα to label OPC (magenta); MYRF to label OLs (yellow). Right: merged image. Scale bar = 25 µm. Bar plots showing mean density by group with standard error for B) for OPC density (cells/mm^2^) C) OL density (cells/mm^2^), and D) OL/OPC ratio. No significant main effect of group was determined from ANCOVA analysis for any of the three density characteristics: OPC density (cells/mm^2^), p = 0.32; OL density (cells/mm^2^), p = 0.54; OL/OPC ratio, p = 0.99. CTRL: control, DS: depressed suicides, DS-CA: depressed suicides with a history of child abuse. OPC: oligodendrocyte precursor cells. OL: oligodendrocytes.

### 3.3 Myelin Area Fraction Coverage

Figure 5a presents a confocal image identifying myelin with MBP and axons with NF-H. Subject information for the density analyses can be found in Supplementary Table 4. The average M2 coefficient across all images was 0.335 ± 0.0162, indicating an average of 33.5% of total BLA axon area covered by myelin. Myelin coverage ranged between subjects from a mean of 0.142 to 0.468 (Fig 5b). No statistically significant differences were observed in M2 coefficient across group (p = 0.075) or any other model factor. However, descriptively, the M2 coefficient was highest in CTRL, followed by DS-CA and lowest in DS. There was no statistically significant correlation between age and M2 coefficient (Supplementary Figure 6d, p=0.57).

**Figure 5.**
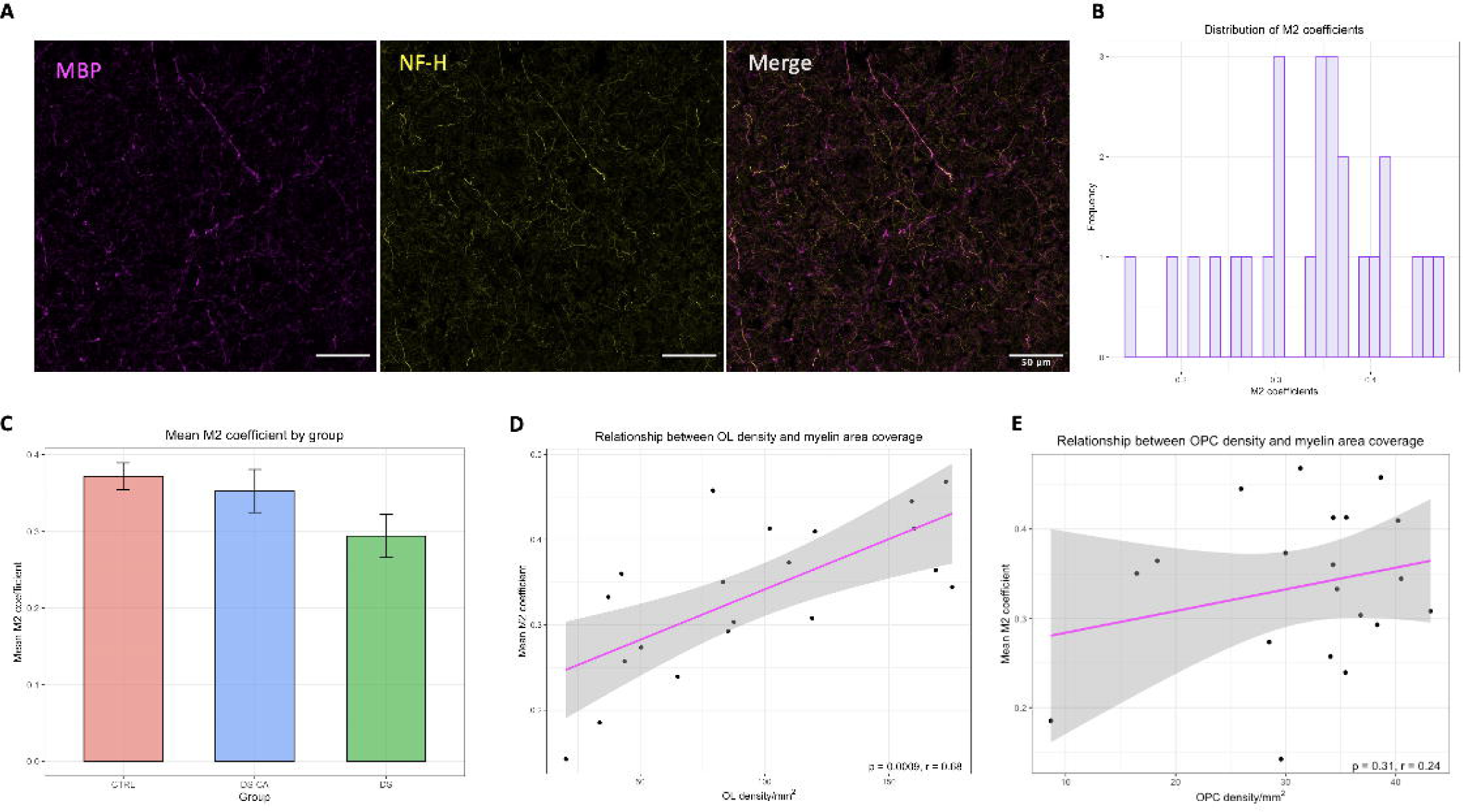
Myelin area fraction coverage analysis. A) Example IF image identifying myelin with MBP (magenta) and axons with NF-H (yellow). Scale bar = 50 *µm.* B) Histogram showing distribution of axonal myelin coverage (M2) determined by colocalization analysis across subjects, showing a wide range of myelin coverage values. C) Bar plot showing mean M2 coefficient by group with standard error. D) Scatter plot with linear trend line showing positive and statistically significant Pearson correlation between M2 and OL density. E) Scatter plot with linear trend line showing positive and non-statistically significant Pearson correlation between M2 and OPC density. IF: immunofluorescence, M2: Mander’s 2 coefficient, OPC: oligodendrocyte precursor cells, OL: oligodendrocytes.

When we considered the overlap of the subjects between the RNAScope cohort and the immunofluorescence cohort (n = 20), we found that mean M2 coefficient showed a strong, statistically significant correlation with OL density (p = 0.0009, r = 0.68), and non-significant positive correlation with OPC density (p = 0.31, r = 0.24) (Figure 5 d,e).

## 4. Discussion

In this study, we developed and validated a novel approach for reliably isolating OPC and OL nuclei via FANS from archived frozen postmortem human brain gray matter. We then applied this new method to perform a targeted BLA gene expression study examining whether there were differences related to depression and/or CA in the BLA. Finally, we performed OPC and OL density measurements and myelin coverage analyses in the BLA comparing these findings between groups. We present this new FANS-based isolation method for frozen human grey matter and provide a characterization of OPC, OL, and myelin in the human BLA as a fundamental contribution to our knowledge of the human amygdala as well as a methodological foundation for future studies on this (and likely other) brain region. Indeed, to our knowledge, there has been no reliable FANS-based method for sorting archived postmortem brain tissue into OPC and OL nuclei, which can then be used for downstream applications. Most methods that incorporate flow cytometry typically deploy cell surface or cytoplasmic markers or a full lineage transcription factor (SOX10 only or OLIG2 only), which does not allow for the separation of OPCs from OLs (Nott et al., 2021; S S Policicchio, 2020). Our combinatorial antibody approach now allows for such separation in a reliable and validated manner. To make better use of archived postmortem human brain tissue in cell type specific studies, it is key to develop protocols that are suitable for nuclei (Wiseman et al., 2023).

In order to characterize the OL-lineage cell population and the extent of myelination in the human BLA comprehensively, we expanded this pipeline to include RNAScope and IF methods to measure OPC/OL cell density and myelin area coverage, respectively. To our surprise, data on human BLA OL-lineage density were challenging to find in the extant literature. The average OPC density we measured was 31.60 ± 1.40 cells/mm^2^, which is consistent with estimates in other human gray matter regions. OPC density in the human vmPFC was found to range between ∼20 and 80 OPCs/mm² (Sibille et al., 2009) and in human cortical grey matter to be around 56 cells/mm² (Strijbis et al., 2017). Human white matter OPC density is considerably higher with estimates ranging from 121 to ∼200 cells/mm² (Ahmed et al., 2013; Strijbis et al., 2017).

Our average overall OL density was 111.31 ± 13.77 cells/mm^2^, compared with an average of 74 cells/mm² in non-specific cortical grey matter and 554 cells/mm² in white matter (Strijbis et al., 2017). Thus, our estimates are broadly consistent with other gray matter regions, although the mean OL density reported in this study is higher than what Strijbis and colleagues found. Finally, we measured an average OL/OPC density ratio of 3.78 ± 0.51, indicating that in the BLA, there is 1 OPC for every 4 OLs. A stereological analysis in mice found 1 OPC for every 7 OLs in the corpus callosum and 1 OPC for every 1 OL in neocortex (Boulanger and Messier, 2017). This difference is likely attributable to region, species and/or methodology (e.g. tissue dissociation methods). Interestingly, our results in the human amygdala are similar to those of a previous snRNAseq study in the adult human amygdala, which showed that OLs comprise 54.4% of all glia while OPCs comprise 11.5% of all glia, indicating an OL/OPC ratio of 4.73 (Tran et al., 2021).

Our quantification of axonal myelin coverage of the BLA yielded a mean of 33.5%, ranging between 14.2% and 46.8% coverage. Our study reports the overall axonal area that is colocalized with myelin – this is not the same as the number or proportion of axons that are myelinated. Future studies could use tracing methods in thicker (40-50 µm) tissue sections to map out the specific neuron types that are myelinated and to what extent. As expected, the M2 coefficient showed a strong correlation with the density of OLs, indicating myelin area fraction coverage. This is congruous with recent work which suggests that number of OL cell bodies can predict the quantity of axonal territory covered by myelin (Xu et al., 2024).

Notably, we made use of our cell type specific gene expression method and histology to investigate whether depressed suicides with or without a history of CA present changes in OPC and/or OL gene expression compared to controls. With respect to group differences, only MOBP in the OL fraction was found to be downregulated in DS-CA, and as such we could not replicate the global downregulation of OL/myelin genes found in the BLA of chronic social stress vs. control mice previously reported (Cathomas et al., 2019). Furthermore, there appears to be no specific impact of CA our histology measures. Given the small sample size of this study, it is possible that we had insufficient power to detect statistically significant effects, so replicating this study in a larger cohort is warranted. It is worth noting that Cathomas and colleagues found that the detected transcriptomic downregulation in the key myelin genes MBP, PLP and CNP, was not reflected at the protein level in western blots, nor did they find a difference in the number of OLs between chronically socially stressed and controls mice (Cathomas et al., 2019). Their BLA dissections contained both white matter and grey matter, and the authors hypothesized that they may observe group differences at the protein level if they would isolate the myelin specifically, rather than measuring at a global level. Our dissections aimed to exclude white matter, possibly explaining the discordant results. It is also worth noting that the specific rodent stressor under study might affect the correspondence of results between rodents and humans. For example, in a translational study by Sibille and colleagues they found a common molecular signature between mice who underwent unpredictable chronic mild stress and a subset of human males with familial depression (Sibille et al., 2009), in which several OL genes were downregulated.

We observed a trend towards lower OL density and lower myelin coverage in both depressed groups (DS-CA and DS) compared to controls. Post-mortem human brain studies that have compared amygdala OL density in donors with major depression with psychiatrically healthy controls are inconsistent. One study found a 47% reduction in OL density in the left amygdala (Strijbis et al., 2017), while another study reported no difference in OL density, though no lateralisation-specific analysis was reported (Williams et al., 2013). Many, though not all, neuroimaging studies of depression and early life adversity report findings that are specific to the right amygdala, particularly with respect to functional connectivity, either in resting state or in response to a task (Damborská et al., 2020; Gee et al., 2013; Ramasubbu et al., 2014). Molecular studies of an ELS rat model have also demonstrated morphological and functional changes affecting synaptic organization and plasticity to be specific to the right BLA (Guadagno et al., 2020, 2018). These hemisphere-specific findings suggest that we should replicate this study using tissue from the right hemisphere, as depression- or CA-related effects might be pronounced on right side. If left side changes are more nuanced as compared to right side, more statistical power might be required to observe group effects in left BLA samples.

Another key finding that emerged in this study concerns age-related changes in BLA gene expression. Aging studies in rodents have demonstrated that each cell type has its own unique transcriptomic program of aging, rather than simply a universal signature of aging, highlighting the importance of cell type specific methods (Allen et al., 2023; Ximerakis et al., 2019). We see several genes in both OPC and OL fractions that have significant relationships with age. This finding is consistent with the well documented observation that especially in human glia, networks of genes change with age (Soreq et al., 2017).

Despite its novelty, this study is not without limitations, with sample size being the foremost among them. The limited sample size means that our statistical analyses might have been underpowered. Furthermore, consistent with 2-3 fold higher rate of suicide in males compared to females (Turecki et al., 2019), the sample under study contains fewer female subjects, making it difficult to robustly detect sex-specific effects in our population. Repeating this OL-lineage/myelin-based analysis on the right BLA, where depression- or CA-associated changes may be more pronounced, could be informative. Furthermore, it would be interesting to repeat these experiments in the central amygdala to explore potential differences with the BLA that could be related to the specific functions played by these amygdalar nuclei. Moreover, the fact that this is a targeted gene expression study, with genes *a priori* selected to investigate OL-lineage development and myelin, constitutes another limitation. An untargeted, whole transcriptome method like RNA sequencing would be useful to understand whether any biologically relevant gene networks are differentially expressed in these groups. There are increasingly crucial roles being attributed to OL-lineage cells beyond their capacity to myelinate. OLs are also central in providing metabolic support to axons (Lee et al., 2012), and OPCs have been recently implicated in axon remodelling and circuit refinement, among other functions (Auguste et al., 2022; Xiao et al., 2022). Lastly, our targeted custom-designed Nanostring codeset only captured exon spanning transcripts in the nucleus. While we believe this to be representative of the biological activity of the cell, there are many transcripts from the nucleus and the cell body that were not examined (Bakken et al., 2018).

In summary, this study on the human BLA has provided 1) a novel method for FANS-based isolation of OPCs and OLs from archived frozen postmortem human brain tissue, 2) baseline OPC and OL densities as well as myelin area coverage estimates, 3) group analyses demonstrating no robust pattern of gene expression differences associated with depression or CA, and 4) a characterization of age-associated changes which are pronounced in both OPC and OL. We anticipate that the methods and results presented here will help to guide future studies on the human amygdala in healthy individuals and patients with brain disorders.

## Data availability statement

Nanostring raw data is available at the following OSF repository: https://osf.io/cx8ya/?view_only=cf95de14c69344eea8ced56429f399fc. The design for the Nanostring probeset, including isoforms can be found as Supplementary File 1. The Fiji script for automatically segmenting the confocal images can be found as Supplementary File 2.

## Acknowledgements

We are deeply indebted to the next of kin who consented to donating the brains of their loved ones. We would like to thank the Nanostring Gene Expression Profiling core facility at the Lady Davis Institute at the Jewish General Hospital. We would also like to thank the staff of the Douglas-Bell Canada Brain Bank and Molecular and Cellular Microscopy Platform at the Douglas Health Research Center for their expert help.

## Author contributions

K.P – Conceptualization, formal analysis, investigation, methodology, project administration, visualization, writing-original draft. E.C. – investigation, methodology, formal analysis, writing – original draft. S.B-B –conceptualization, methodology. M-A.D. – methodology, Writing – review and editing. V.Y - methodology, investigation, writing – review and editing. R.M – investigation, methodology, resources, writing – review and editing. A.I.P. – conceptualization, project administration. methodology, resources, writing – review and editing. A.S. – resources, writing – review and editing. C.R.P. – conceptualization, writing – review and editing, G.T. – resources. N.M. – conceptualization, project administration, funding acquisition, resources, supervision, writing – review and editing.

