## Supplementary File 2 for "Characterizing Oligodendrocyte-Lineage Cells and Myelination in the Basolateral Amygdala: Insights from a Novel Methodology in Postmortem Human Brain"

// Script created with assistance from ChatGPT

// Define the directory containing the image files 
dir = "";

// Get a list of all files in the directory
list = getFileList(dir);

// Loop through each file in the directory
for (i = 0; i < lengthOf(list); i++) {
    // Check if the file is an actual .oif file and not an auxiliary file
    if (endsWith(list[i], ".oif")) {
        // Open the current .oif image file
        open(dir + list[i]);

        // Check if the image size matches the expected dimensions (1600x1600)
        if (getWidth() == 1600 && getHeight() == 1600) {
            // Perform max intensity projection while retaining all channels
            run("Z Project...", "projection=[Max Intensity]");
            
            myTitle = getTitle();  // Get the title after projection

                // Bring up brightness and contrast windows
                run("Brightness/Contrast...");
                run("Channels Tool...");
                run("Split Channels");

                // Make otsu binary mask of channel 2
                selectWindow("C2-" + myTitle);
                setAutoThreshold("Otsu dark");
                setOption("BlackBackground", false);
                run("Convert to Mask");
                // Save the binary mask of channel 2
                saveAs("tiff", dir + File.nameWithoutExtension + "_otsu_bin_Channel2.tif");
                if (isOpen("Save As")) {
                    close("Save As");
                    print("Error saving binary mask of channel 2 for file: " + list[i]);
                } else {
                    print("Binary mask of channel 2 saved successfully for file: " + list[i]);
                }

                // Make otsu binary mask of channel 3
                selectWindow("C3-" + myTitle);
                setAutoThreshold("Otsu dark");
                setOption("BlackBackground", false);
                run("Convert to Mask");
                // Save the binary mask of channel 3
                saveAs("tiff", dir + File.nameWithoutExtension + "_otsu_bin_Channel3.tif");
                if (isOpen("Save As")) {
                    close("Save As");
                    print("Error saving binary mask of channel 3 for file: " + list[i]);
                } else {
                    print("Binary mask of channel 3 saved successfully for file: " + list[i]);
                }
            
        } else {
            print("Skipping file due to size mismatch: " + list[i] + " (" + getWidth() + "x" + getHeight() + ")");
        }

        // Close the current image
        close();
    } else {
        print("Skipping non-oif file: " + list[i]);
    }
}
