## Supplementary Material for "Characterizing Oligodendrocyte-Lineage Cells and Myelination in the Basolateral Amygdala: Insights from a Novel Methodology in Postmortem Human Brain"

**Supplementary Tables**

Supplementary Table 1 - Subject information for Nanostring gene expression analysis, for subjects for which there was useable *OL fraction* results. Data are displayed as mean ± standard error of the mean. P-values are derived from one-way anovas.

|  | **CTRL** | **DS-CA** | **DS** |
| --- | --- | --- | --- |
| **N total** | 17 | 15 | 18 |
| **N female** | 2 | 2 | 2 |
| **Age** (p=0.81) | 49.35 ± 2.98 | 46.50 ± 3.31 | 47.65 ± 3.14 |
| **PMI** (p=0.55) | 49.08 ± 5.14 | 53.96 ± 3.38 | 55.92 ± 4.77 |
| **pH** (p=0.40) | 6.24 ± 0.085 | 6.36 ± 0.075 | 6.37 ± 0.068 |
| **Known antidepressant** | 0 | 7 | 10 |
| **Mean SOX10+%** (P=0.99) | 31.44 ± 2.58 | 31.18 ± 2.84 | 31.07 ± 2.12 |

Supplementary Table 2 - Subject information for Nanostring gene expression analysis, for subjects for which there was useable *OPC fraction* results. Data are displayed as mean ± standard error of the mean. P-values are derived from one-way anovas.

|  | **CTRL** | **DS-CA** | **DS** |
| --- | --- | --- | --- |
| **N total** | 16 | 16 | 14 |
| **N female** | 2 | 2 | 2 |
| **Age** (p=0.93) | 48.94 ± 3.14 | 47.56 ± 3.32 | 49.14 ± 3.48 |
| **PMI** (p=0.33) | 47.40 ± 5.17 | 55.58 ± 3.69 | 56.76 ± 5.45 |
| **pH** (p=0.46) | 6.23 ± 0.090 | 6.36 ± 0.075 | 6.36 ± 0.080 |
| **Known antidepressant** | 0 | 7 | 9 |
| **Mean SOX10+%** (P=0.99) | 31.91 ± 2.70 | 31.85 ± 2.72 | 31.47 ± 2.13 |

Supplementary Table 3 – Subject information for RNAScope density analysis. Data are displayed as mean ± standard error of the mean. P-values are derived from one-way anovas. P-values < 0.05 are italicized.

|  | **CTRL** | **DS-CA** | **DS** |
| --- | --- | --- | --- |
| **N total** | 10 | 9 | 10 |
| **N female** | 4 | 2 | 2 |
| **Age** (p=0.78) | 45.60 ± 5.52 | 48.56 ± 3.06 | 44.10 ± 4.29 |
| **PMI** (p=0.67) | 57.04 ± 5.71 | 50.03 ± 6.08 | 52.05 ± 5.26 |
| **pH** *(p=0.041)* | 6.187 ± 0.082 | 6.439 ± 0.065 | 6.367 ± 0.058 |
| **Known antidepressant** | 1 | 7 | 4 |

Supplementary Table 4 – Subject information for immunofluorescence myelin coverage analysis. Data are displayed as mean ± standard error of the mean. P-values are derived from one-way anovas.

|  | **CTRL** | **DS-CA** | **DS** |
| --- | --- | --- | --- |
| **N total** | 7 | 9 | 10 |
| **N female** | 2 | 2 | 4 |
| **Age** (p=0.55) | 52.43 ± 5.79 | 50.11 ± 2.14 | 46.50 ± 3.35 |
| **PMI** (p=0.50) | 56.06 ± 6.22 | 46.54 ± 5.28 | 48.35 ± 5.50 |
| **pH** (p=0.084) | 6.21 ± 0.099 | 6.45 ± 0.070 | 6.424 ± 0.065 |
| **Known antidepressant** | 0 | 5 | 6 |

**Supplementary Figures**

**A**

**B**

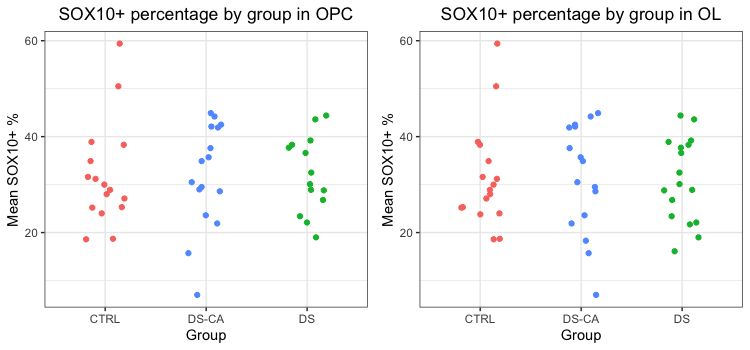

**Supplementary Figure 1 –Distribution of SOX10+ percentage across** **groups**. Scatter plot demonstrating mean SOX10+ percentage in each FANS sort by group, for A) all subjects for which there were usable OPC fractions, and B) all subjects for which there were useable OL fractions. There is no statistically significant difference in either A (p = 0.99) or B (p = 0.99) indicating that differences in gating are unlikely to be driving the gene expression results.

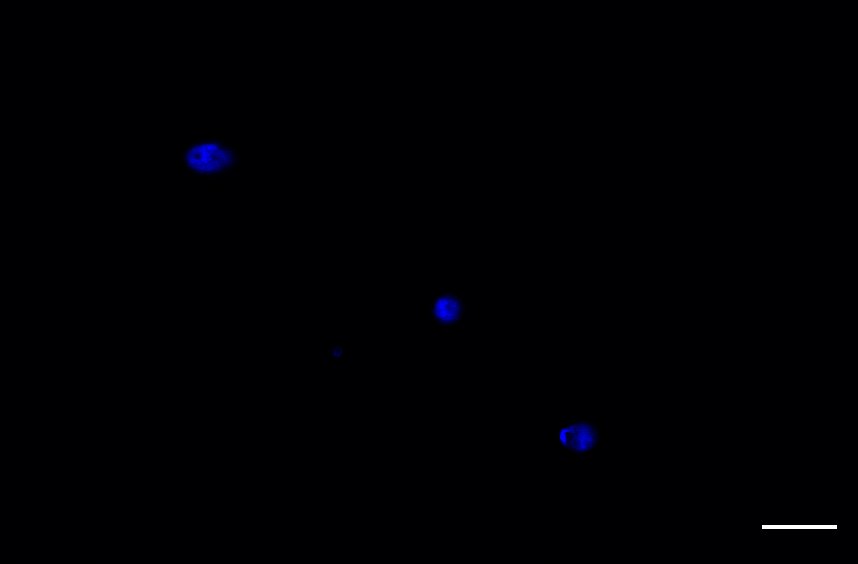

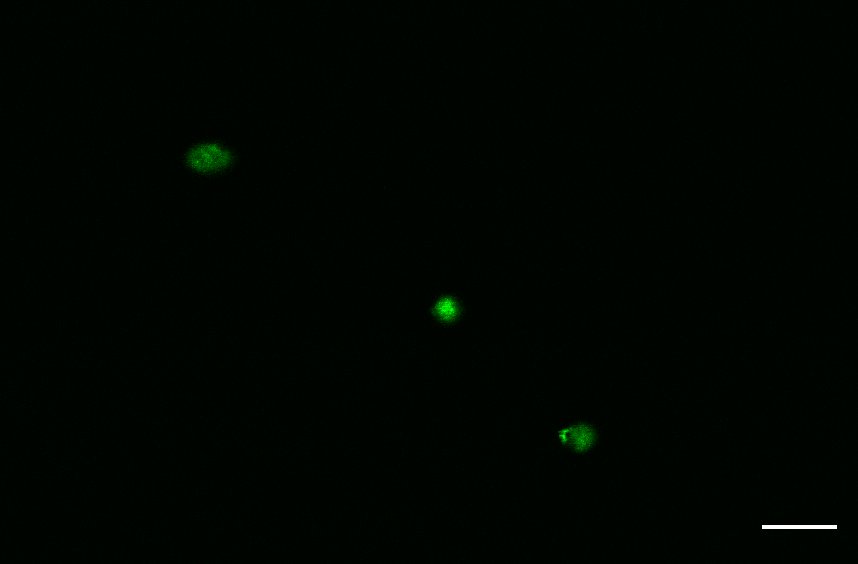

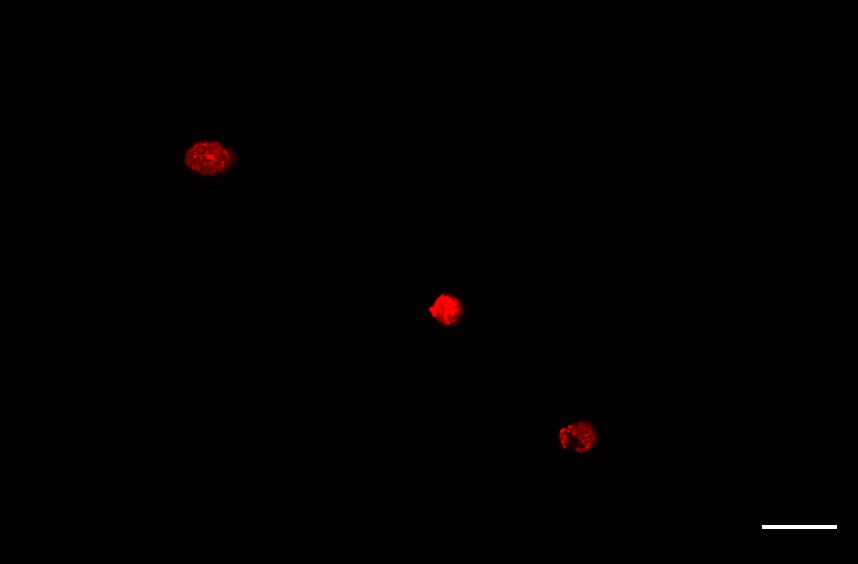

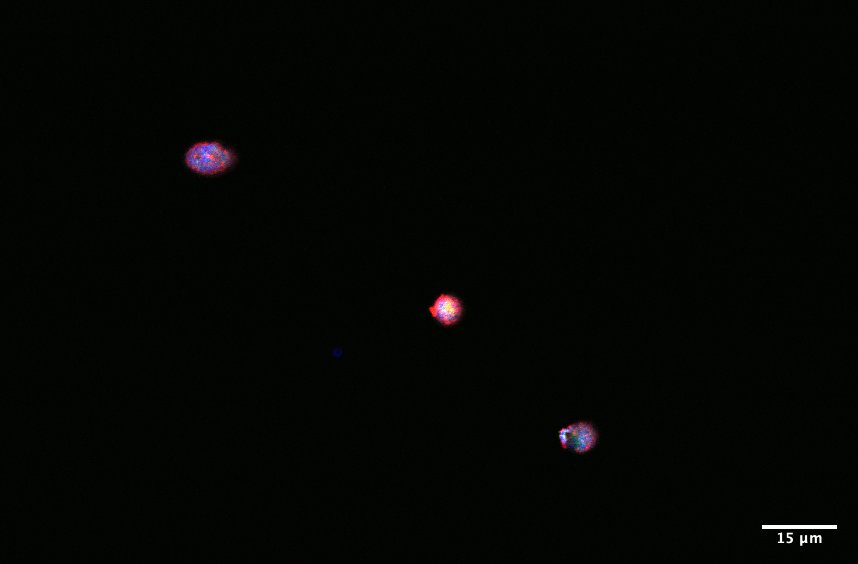

DAPI

SOX10

CRYAB

Merge

**Supplementary Figure 2 – Micrograph of SOX10+/CRYAB+ nuclei.** SOX10 is a pan-OL-lineage marker. SOX10+/CRYAB+ is labels OLs and SOX10+/CRYAB- labels OPCs. The above micrograph shows an example of SOX10+/CRYAB+ nuclei at 40x magnification. Scale bar = 15 μm.

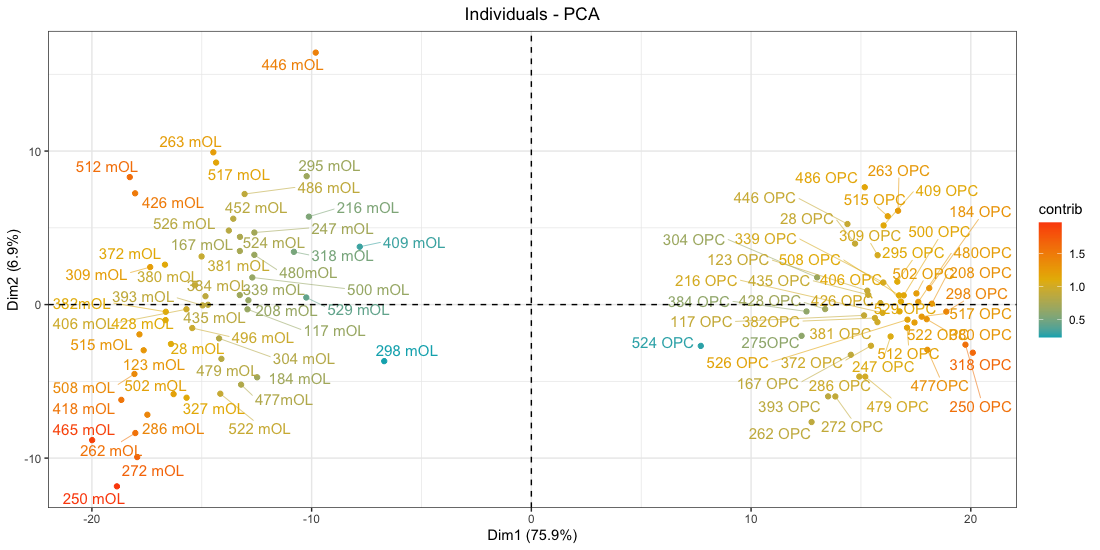

**Supplementary Figure 3 – OL and OPC samples separated along first principal component.** PCA plot showing that along the first principal component (dim1) which accounts for 75.9% of the variance within the data, OL and OPC samples show a clear separation. OL samples show negative values for dim1, while OPC samples show positive values for dim1. The color bar shows the contribution of each sample to the principal component, with red colors indicating a higher contribution and blue colors indicating a lower contribution.

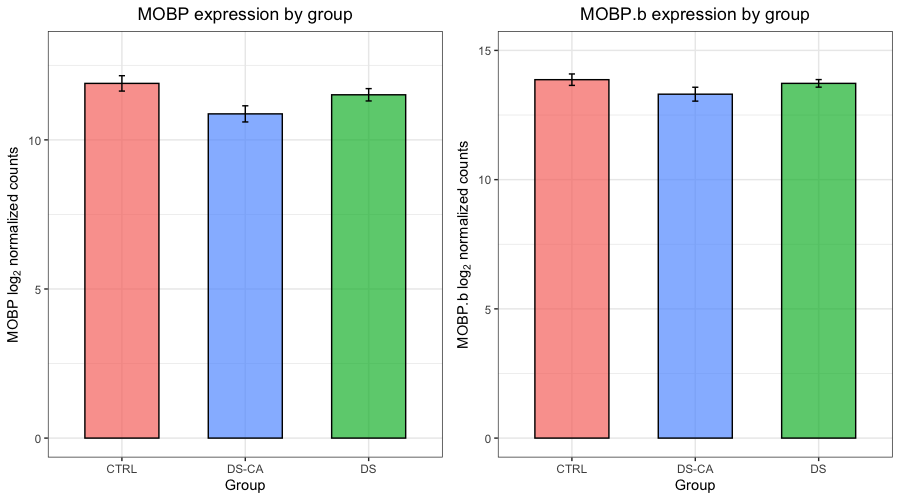

**A**

**B**

*****

**Supplementary Figure 4 – MOBP expression by group.** Two probes of MOBP were created for the Nanostring codeset, in order to span all important isoforms of the gene. Mean expression across groups of A) MOBP and B) MOBP are demonstrated in the bar plots. The standard error is represented with black error bars. Only A) MOBP shows a statistically significant difference between groups (BH corrected p-value = 0.044), though the overall pattern of expression is the same in both probes.

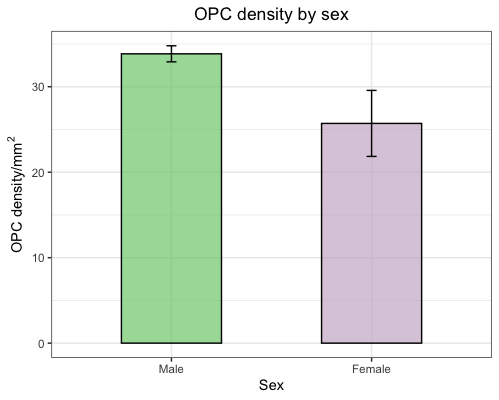

**Supplementary Figure 5- OPC density is lower in females**. Bar plot showing OPC density as a function of sex, displaying a statistically significant reduction in females compared to males (p = 0.0057).

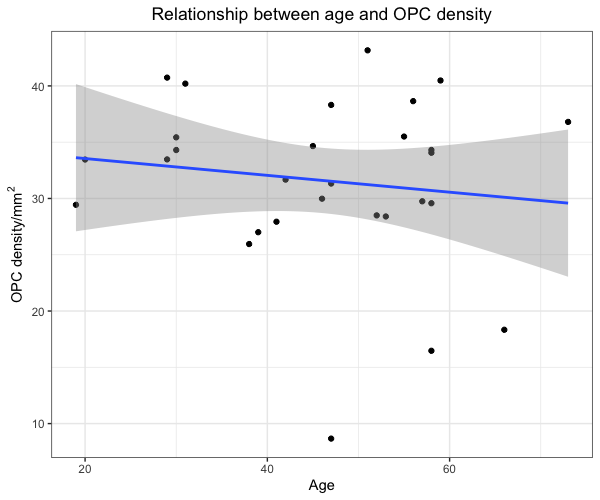

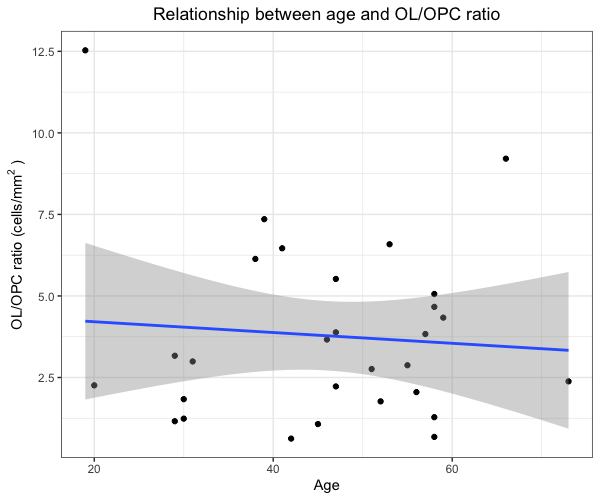

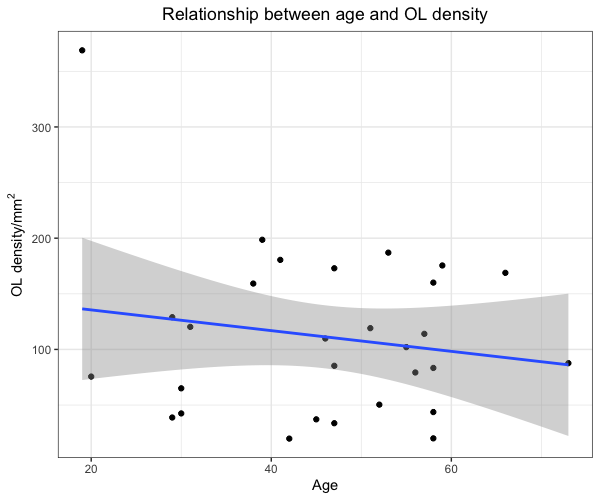

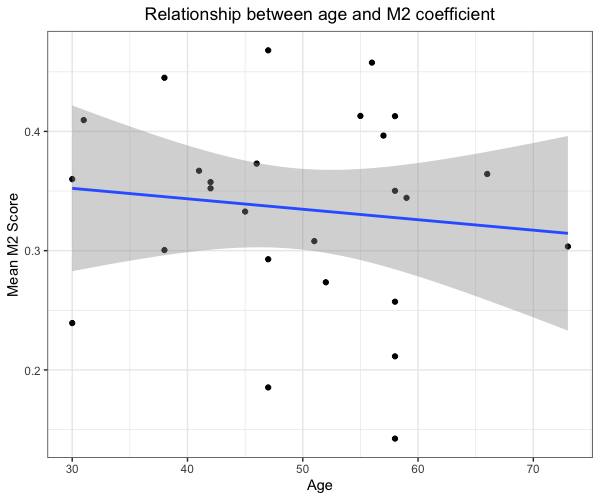

**A**

**B**

**C**

**D**

**Supplementary Figure 6 – Relationships between age and histology measures**. No statistically significant relationship was found between and a) OPC density (p = 0.49, b) OL density (p = 0.38), and c) OL/OPC ratio (p = 0.67)., and d) M2 coefficient (p = 0.57)
